# Stability and Processing Impacts on Faecal Microbiota Transplant Products: An Integrated Metagenomic-Culturomic Analysis

**DOI:** 10.1101/2025.09.02.673862

**Authors:** Calum J. Walsh, Taylor Harshegyi-Hand, Louise M. Judd, Khin Chaw, Elizabeth Connolly, Iain B. Gosbell, Jason C. Kwong, Benjamin P. Howden, Timothy P. Stinear

## Abstract

**Background:** Faecal microbiota transplantation (FMT) is an increasingly used microbiome-based therapy, whose clinical success depends on the stability and viability of the donor microbiota. Few studies have systematically evaluated these parameters for FMT preparations. Here, we integrate metagenomic sequencing with culturing-based recovery of viable microbes to comprehensively characterise FMT preparations from healthy Australian donors.

**Results:** Metagenomic profiling of bacterial communities from repeat FMT faecal donations revealed high temporal stability within individual donors, supporting donor suitability for consistent and standardised FMT production. The community structure and diversity of final FMT products remained largely consistent with their source material, reflecting limited disruption during processing. Culturing and sweep metagenomics showed bacterial species representing 98% of faecal microbiota abundance was recoverable, underscoring the potential of FMTs to deliver a viable and functionally active bacterial community to a recipient. When FMT donor microbiomes were compared to >7,000 healthy gut microbiomes from 27 countries, donor samples were distinct but clustered most closely with other Westernised populations, while also exhibiting significantly higher diversity than most global cohorts. These results highlight the geographic specificity of gut microbiomes and the need to consider population context in donor selection.

**Conclusions:** This study strengthens the evidence base for FMT product standardisation and viability, with implications for FMT clinical use, donor screening protocols, and associated regulatory frameworks.

## Introduction

Faecal microbiota transplantation (FMT) has emerged as a highly effective treatment for recurrent *Clostridioides difficile* infection (CDI)^1^, with growing interest in its application for other microbiome-associated conditions, including inflammatory bowel disease^2^, hepatic encephalopathy^3^, autism^4^, fibromyalgia^5^, melanoma^6^, acute graft-versus-host disease^7^, and colonisation with multidrug-resistant organisms (MDROs)^8^. By introducing a diverse and functionally competent microbial community, FMT aims to restore key ecosystem functions of the gut microbiome that have been disrupted by antibiotics or disease, irrespective of whether that disruption contributes to (or causes) disease development. Recent clinical trials and cohort studies have expanded the evidence base for FMT beyond CDI, exploring FMT potential in modulating host immunity^9^, metabolic syndrome^10^, and colonisation resistance^11^.

As clinical applications broaden, there is increasing interest in defining not only what makes an optimal FMT donor, but also what constitutes an optimal donation. Optimal donors are typically considered to be individuals in good overall health who lack factors that could alter the “normal” structure or function of the gut microbiome, such as chronic disease or recent exposure to antimicrobials.^12^. In contrast, optimal donations refer to specific stool samples that possess favourable microbiological characteristics – such as high microbial diversity, the presence of key species or strains, and an absence of potentially harmful taxa. While donor screening has traditionally focused on overall health status and ensuring samples are free of pathogenic organisms, emerging evidence suggests that both donor attributes and donation-specific features can influence treatment outcomes^12,13^. A key question is whether selecting donors with highly consistent faecal microbiota profiles, and ensuring stability across repeated donations, can improve the reproducibility and predictability of FMT products beyond administration for CDIs. This, in addition to understanding the compositional fidelity of FMT preparations relative to their source material and the extent to which viable microbes are retained during processing, is essential for developing reliable, standardised FMT products, and discerning the microbial traits that influence treatment outcomes - a key step towards the rational design of live biotherapeutic products.

While the gut microbiome is known to be highly individualised and temporally stable in healthy individuals^14–17^, there is limited evidence on the consistency of donor microbiome profiles over time in the context of FMT. This is particularly relevant for so-called “super-donors” - individuals whose FMTs yield significantly improved patient outcomes^18^ – who are likely to be relied upon for repeated donations over time, making intra-donor microbiome stability a key factor in maintaining consistent therapeutic efficacy.

Additionally, although FMT aims to transfer a representative and viable microbial community, the extent to which viable microbes are retained in processed FMT preparations remains poorly characterised. Beyond viability, another important dimension of FMT efficacy relates to the representativeness of donor microbiomes within a global context. A further gap lies in understanding how the microbiomes of FMT donors, particularly within the Australian population, compare to those from other global cohorts. Given that microbiome composition can be influenced by host factors such as ethnicity, age, geography, and lifestyle, it is important to contextualise donor microbiomes to assess their representativeness and potential generalisability across populations.

To address key questions around the composition, consistency, and viability of FMTs, we employed an integrated approach combining metagenomics and culturomics. First, we assessed the stability of the microbiome in repeat donations collected over time from prospective donors to determine intra-donor consistency. We then compared the microbiome profiles of source faecal samples to their derived FMT products, evaluating the fidelity of microbial composition and diversity across these paired samples. To assess the viability of microbial populations post-processing, we applied sweep metagenomics to quantify the proportion of the original microbial community retained in FMTs. Finally, we contextualised these donor microbiomes by comparing them to healthy gut microbiomes from diverse global populations. Together, these analyses provide a detailed characterisation of FMT composition and quality, offering insights relevant to both clinical use and microbiome-based therapeutic development.

## Methods

### Donor Recruitment and Screening

Stool samples and FMT products used in this study were collected as part of Australian Red Cross Lifeblood’s ongoing microbiome program. Prospective FMT donors initially completed an electronic eligibility quiz to determine whether they met the basic criteria to proceed. Those who passed this initial screen underwent a comprehensive assessment that included detailed questionnaires, a physical examination, and the collection of blood, stool, and deep nasal/throat swabs for laboratory screening. Individuals who meet all eligibility and screening requirements were then invited to provide stool donations at a good manufacturing practice-compliant microbiome facility. Their first donation had to occur within 30 days of completing the initial screening tests.

Donors attended the facility to provide stool samples multiple times over a two-week production cycle, with the first day of this cycle designated as “day 0”. Prior to each donation, donors completed a brief re-assessment questionnaire to monitor for any changes in health status. At the conclusion of the cycle, “bookend” testing - including repeat blood, stool, and deep nasal/throat swabs - was conducted between 21 and 25 days after the ‘day 0’ donation. The results of this follow-up testing determined the final release of the FMT products derived from that donation cycle.

Temporal stability of repeat donations was assessed using samples collected at these three defined timepoints: Timepoint A (initial screening, up to 30 days before donation), Timepoint B (during the donation cycle, days 0-14), and Timepoint C (bookend testing, days 21-25).

### FMT product processing

Donated stool was assessed visually for suitability of processing by trained technicians. Suitable donations were homogenised with 0.9% sterile saline and filtered to remove particles > 1mm. Glycerolyte 57 (Fisher Scientific) was added as cryoprotectant at 10% concentration of the FMT product.

### Preparation of samples for metagenomics and culturomics

Faecal samples were stored in DNA/RNA Shield (Zymo Research) to preserve nucleic acids and microbiome structure, but the inactivation of microbes rendered them unsuitable for sweep metagenomics. All downstream handling and preparation of samples for sequencing was performed by the Centre for Pathogen Genomics Innovation Hub (CPG-IH) at the University of Melbourne. FMT sweep samples were generated by serially diluting and plating FMT products in duplicate on YCFA agar from 10^-4^ to 10^-12^ before culturing at 37°C under anaerobic conditions for 72 hours. Twenty-four individual colonies per donation were picked from these plates and re-streaked for purity on YCFA agar before being processed for whole genome sequencing. Colonies were selected to maximise morphological diversity, with the aim of capturing a broad range of genetic and taxonomic diversity. The remaining growth from the sweep plates was counted and scraped clean into sterile phosphate buffered saline for metagenomic sequencing. Bacteria were resuspended in 300μl sterile PBS and placed in a PowerbeadPro tube (QIAGEN). Bacteria were lysed by bead beating (Precellys 24 5000 2 x 20 seconds). DNA from isolates or plate sweeps was purified from 50μl of lysed solution using 30μl of Illumina Purification beads (20060059) and eluted in 30μl of Nuclease Free Water. DNA was extracted from the FMT products direct from syringe, and source faecal samples using a bead beating extraction protocol and modified PowerFaecal Pro Kit (QIAGEN) on the EZ2 instrument (QIAGEN). Illumina libraries were prepared using Illumina DNA Prep (20060059) at quarter reaction volumes. Libraries were sequenced on a NextSeq2000 P4 XLEAP-SBS™ Reagent Kit (300 Cycles) (20100992), targeting 20 million 2 × 150 bp read pairs per sample.

### Quality Control of Metagenomic Data

The raw metagenomic sequencing reads were processed using *fastp*^19^ for quality filtering and adapter trimming, with *Bowtie2*^20^ for removal of human-derived reads. Specifically, read pairs were removed if either mate aligned to the human reference genome (GRCh38_noalt_as) to eliminate potential host contamination. Taxonomic profiling was performed at the species level using *Kraken*2^21^ and *Bracken*^22^, with the PlusPF^23^ database as the reference. Residual human data (species classification *Homo sapiens* in Bracken profiles) were removed from the resulting species abundance tables using the *tidyverse*^24^ library (v. 2.0.0) in *R*^25^ (v. 4.5.0), which was also used for downstream data manipulation and visualisation unless otherwise stated. Species abundance tables were filtered to only retain species with a mean relative abundance of at least 0.1% across all samples.

### Community-level Analysis of Metagenomic Data

Microbial diversity within and between samples was assessed using the *vegan*^26^ package (v. 2.6-10) in *R*. Alpha diversity was calculated using the Shannon index, while beta diversity was measured using Bray-Curtis dissimilarity. Beta diversity was visualised using principal coordinates analysis (PCoA).

Statistical comparisons were conducted to evaluate differences in diversity metrics across groups. Non-parametric tests, including the Kruskal-Wallis test and Wilcoxon rank-sum test, were applied using the *rstatix*^27^ package (v. 0.7.2). Where appropriate, mixed-effects linear models were implemented with the *lme4*^28^ package (v. 1.1-37) to account for repeated measures and the *lmerTest*^29^ package (v. 3.1-3) was used to test for statistical significance. Differences in overall microbial community composition were tested using permutational multivariate analysis of variance (PERMANOVA), as implemented in the *vegan* package.

### Species-level Analysis of Metagenomic Data

Species-level differential abundance testing was performed using the *LinDA*^30^ framework employed in the *R* package *MicrobiomeStat*^31^ (v. 1.2), with abundance-filtered species count data as input. For comparisons between sample sources, donation ID was included as a random effect to account for inter-individual variation in microbiome composition. To detect taxonomic clades showing consistent enrichment patterns between sample sources, species names were converted to full taxonomic lineages using *TaxonKit*^32^ and differential abundance log2 fold changes were analysed using *TaxSEA*^33^ with FDR < 0.1 accepted as significant.

Metagenomic relative abundance data generated by *Bracken* was used to evaluate the extent to which viable species from stool and FMT samples were recovered by sweep metagenomics. Species with non-zero relative abundance in the corresponding FMT Sweep sample were considered “recovered” (i.e., viable and culturable), and those absent were considered “not recovered”. This presence/absence designation was used as a binary indicator of viability, avoiding potential biases introduced by differential growth rates or culturing efficiency. To estimate the fraction of the original microbiome composition that was recoverable, two complementary metrics were calculated:

1. Abundance-weighted recovery: For each Donation and Source (Stool or FMT), the total relative abundance of species classified as “Recovered” was calculated. This metric reflects the proportion of the microbial community (by relative abundance) that was recovered as viable isolates.
2. Presence/absence-based recovery: For each Donation and Source (Stool or FMT), the proportion of detected species (i.e., those with relative abundance > 0) that were also recovered in the paired FMT Sweep sample was calculated. This metric captures recovery based on species richness rather than abundance.

Mixed-effects linear models were used to compare recovery between Stool and FMT samples, with Donation included as a random effect. Unrecovered species were identified and summarised to better understand which taxa were consistently missed under the culturing conditions used.

### Strain-level Analysis of Metagenomic Data

Strain-level profiling of metagenomic samples was performed using *inStrain*^34^ with a database constructed from draft genomes of cultured isolates dereplicated at 98% average nucleotide identity (ANI) using *dRep*^35^. Paired-end metagenomic reads were aligned to this database using *Bowtie2*, and profiles were generated with *inStrain profile*.

Strain comparisons were conducted using *inStrain compare*, with strain sharing defined as population ANI ≥ 99.999% and ≥ 50% genome coverage, following the author-recommended cutoffs. Strain clusters identified by *inStrain* were used to assess persistence and distribution across donations and sample sources. Community-level similarities between donations and sample types were calculated by using a presence/absence matrix of strain clusters to calculate binary Jaccard distance which was then visualised by PCoA.

### Quality Control of Isolate Genomic Data

Raw isolate sequencing reads were processed using *fastp* for quality filtering and adapter removal. Draft assembles were generated by *Shovill*^36^, discarding contigs shorter than 1000bp. Taxonomy was assigned to assembled genomes using *GTDB-tk* (v. 2.1.1)^37^. Isolates were excluded from downstream analyses if they did not have a suitable MSA length of marker genes reported by *GTDB-tk*. A phylogenetic tree was constructed from all draft genome assemblies which passed quality control using the *de_novo_wf* functionality in *GTDB-tk* with the *Actinobacteriota* phylum as outgroup.

### Analysis of Isolate Genomic Data

To define species-level clusters within the *Collinsella* genus, complete-linkage hierarchical clustering was performed using pairwise average nucleotide identity (ANI) values computed by *FastANI*^38^. A distance matrix was constructed as (100 – ANI%), and clustering was applied using the *hclust* function in *R*. Clusters were defined using a distance threshold of 5 (corresponding to 95% ANI), based on the species boundary proposed for prokaryotes by the authors of *FastANI*. Cluster membership was extracted using *cutree*. To visualise the resulting clusters, a dendrogram was plotted using the *ggtree*^39^ package, with cluster assignments mapped to the tree as coloured tip points.

To assess whether alignment rates of metagenomic reads to genome assemblies of isolated strains differed between sample sources, we fit a linear mixed-effects model with the percentage of aligned reads as the response variable, Source as a fixed effect, and Donation as a random intercept to account for inter-individual variation. Pairwise comparisons between sources were performed using estimated marginal means via the *emmeans*^40^ package (v. 1.11.1), with Tukey adjustment for multiple testing.

### Comparison of Stool Metagenomic Data to Public Datasets

To compare the faecal microbiome composition of healthy Australian FMT donors to that of healthy individuals from other countries, we re-analysed Stool samples from the FMT donations using *MetaPhlAn3*^41^ with default parameters and the mpa_v30_CHOCOPhlAn_201901 database, ensuring compatibility with taxonomic profiles available in the *curatedMetagenomicData*^42^ package for *R*. A global comparison dataset was constructed by extracting *MetaPhlAn3* profiles from samples that met the following criteria: *body_site == “stool”*, *disease == “healthy”*, *study_condition == “control”*, *age_category == “adult”*, *treatment == “no”* or *NA*, and *number_reads ≥ 10000000*. Only samples with an assigned country were retained. Alpha diversity (Shannon index) and beta diversity (Bray–Curtis dissimilarity) were calculated as described above. Differences in Shannon diversity across countries were assessed using a linear mixed-effects model with country as a fixed effect and subject as a random effect to account for repeated sampling. To evaluate differences in community composition, pairwise Bray-Curtis dissimilarities between Australian samples and those from each other country were modelled using a linear mixed-effects approach. Intra-Australian comparisons served as the reference group, and subject was included as a random effect to account for repeated measures. All comparison samples were included in diversity analyses and statistical comparisons. To improve interpretability in beta diversity ordination, up to 30 representative samples were selected per country based on lowest median intra-country Bray-Curtis dissimilarity, reflecting those most similar to their country’s overall profile. A full list of all samples and the selected representative subsets is provided in the supplementary material.

## Results

### Donor Microbiomes Show High Intra-Individual Consistency Over Time

To determine whether repeated faecal donations from the same donor contained a consistent microbial population, we used donations from 10 donors collected by Australian Red Cross Lifeblood as part of their ongoing donor screening program. Two donors failed the initial screen so only provided one sample. The remaining 8 donors provided either 2 or 3 samples.

Microbiome diversity and composition remained relatively stable across repeat donations from the same individual, with inter-individual differences contributing more to overall variation than temporal changes from each donor (Fig. 1). A linear mixed-effects model including donor as a random effect found no significant change in Shannon diversity between the first and second donations (estimate = 0.067, *p* = 0.395), while a modest but statistically significant decrease was observed by the third donation (estimate = -0.283, *p* = 0.018) (Fig. 1A).

**Figure 1.**
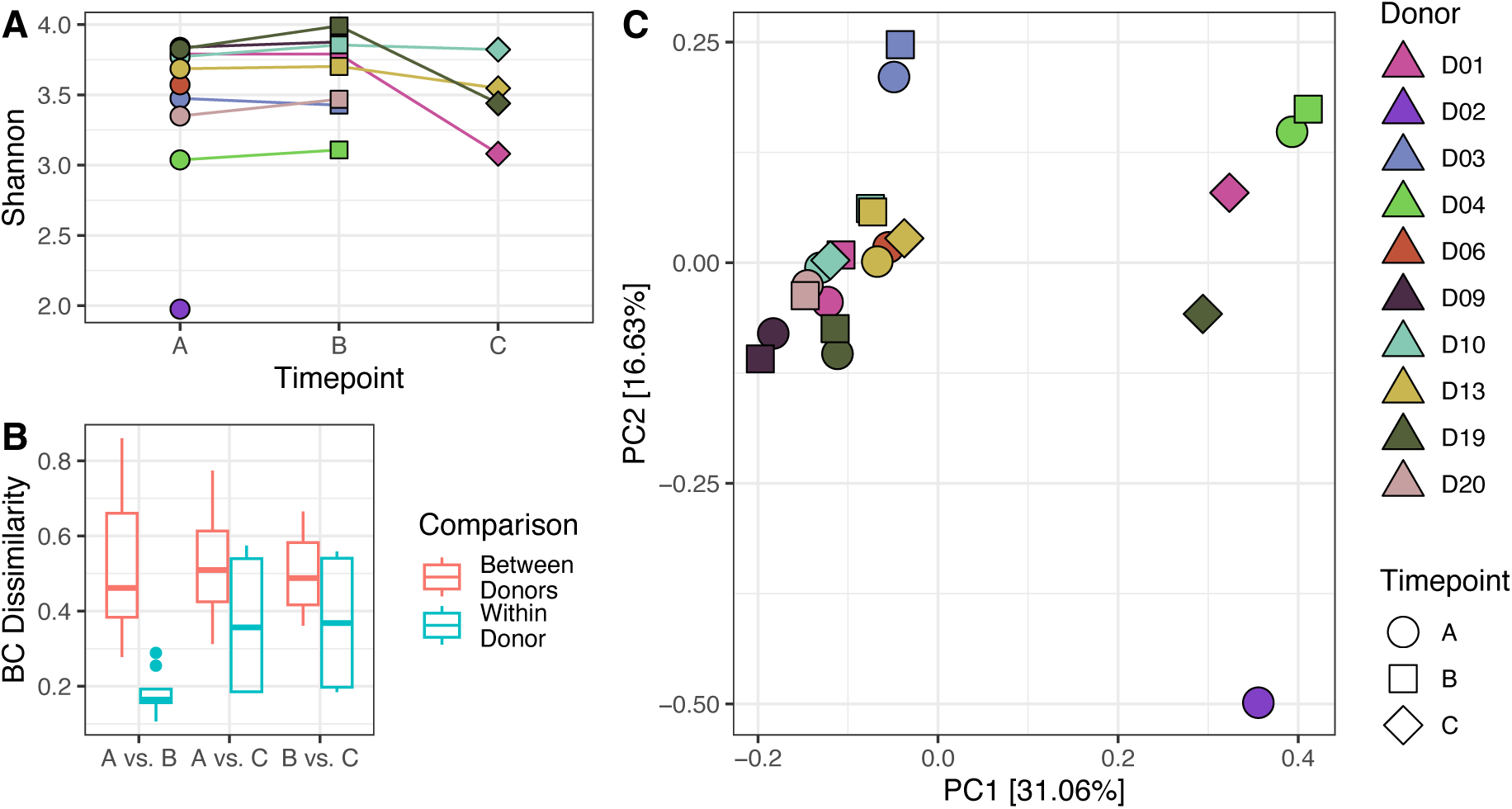
Consistency of donor profiles over time. A) Alpha diversity (Shannon index) trajectory of faecal microbiome samples by Donor and Timepoint. B) Pairwise Bray-Curtis dissimilarities comparing within-donor versus between-donor sample comparisons across each combination of timepoints. C) Principal Coordinate Analysis of Bray-Curtis dissimilarities between samples. For panels A and C, each point represents a single faecal microbiome sample; points are colour-filled by Donor and shaped by Timepoint.

PERMANOVA of Bray-Curtis dissimilarities including both Donor and Timepoint showed that Donor identity accounted for most of the variation in microbiome composition (R² = 0.79, *p* = 0.001), with a smaller but statistically significant contribution from Timepoint (R² = 0.079, *p* = 0.002) (Fig. 1C). When controlling for repeated measures using Donor as a blocking factor, the effect of Timepoint was no longer statistically significant (R² = 0.079, *p* = 0.057), suggesting modest and inconsistent temporal shifts within individuals.

Pairwise Bray-Curtis dissimilarities were significantly lower between samples from the same donor compared to samples from different donors (Two-sample Wilcoxon test, *p* = 6x10^-6^).

There was a significant global difference in Bray-Curtis dissimilarity between timepoint pairs within donors (i.e., A vs B, A vs C, and B vs C; Kruskal-Wallis rank sum test, *p* = 0.042) (Fig. 1B). However, no individual pairwise comparison was statistically significant after Holm adjustment, suggesting that while temporal changes in microbiome composition may occur between donations, these differences are subtle or not consistent across donors.

### Processed FMTs Closely Resemble Source Stool Microbiomes

To assess the impact of FMT processing on the microbial population we used metagenomics and culturomics to compare the microbiome diversity and composition of ten stool samples to their derived FMTs. These stool samples were collected from nine donors, with two (Donation02 and Donation03) originating from the same individual on consecutive days.

A linear mixed-effects model including donation as a random effect indicated that Shannon diversity did not differ significantly between stool and FMT samples (estimate = 0.084, *p* = 0.416). However, FMT Sweep samples (FMT samples subjected to culturomics and sweep metagenomics, see methods) had significantly lower Shannon diversity compared to the source stool (estimate = -0.77, *p* = 4.4x10^-7^) (Fig. 2A).

**Figure 2.**
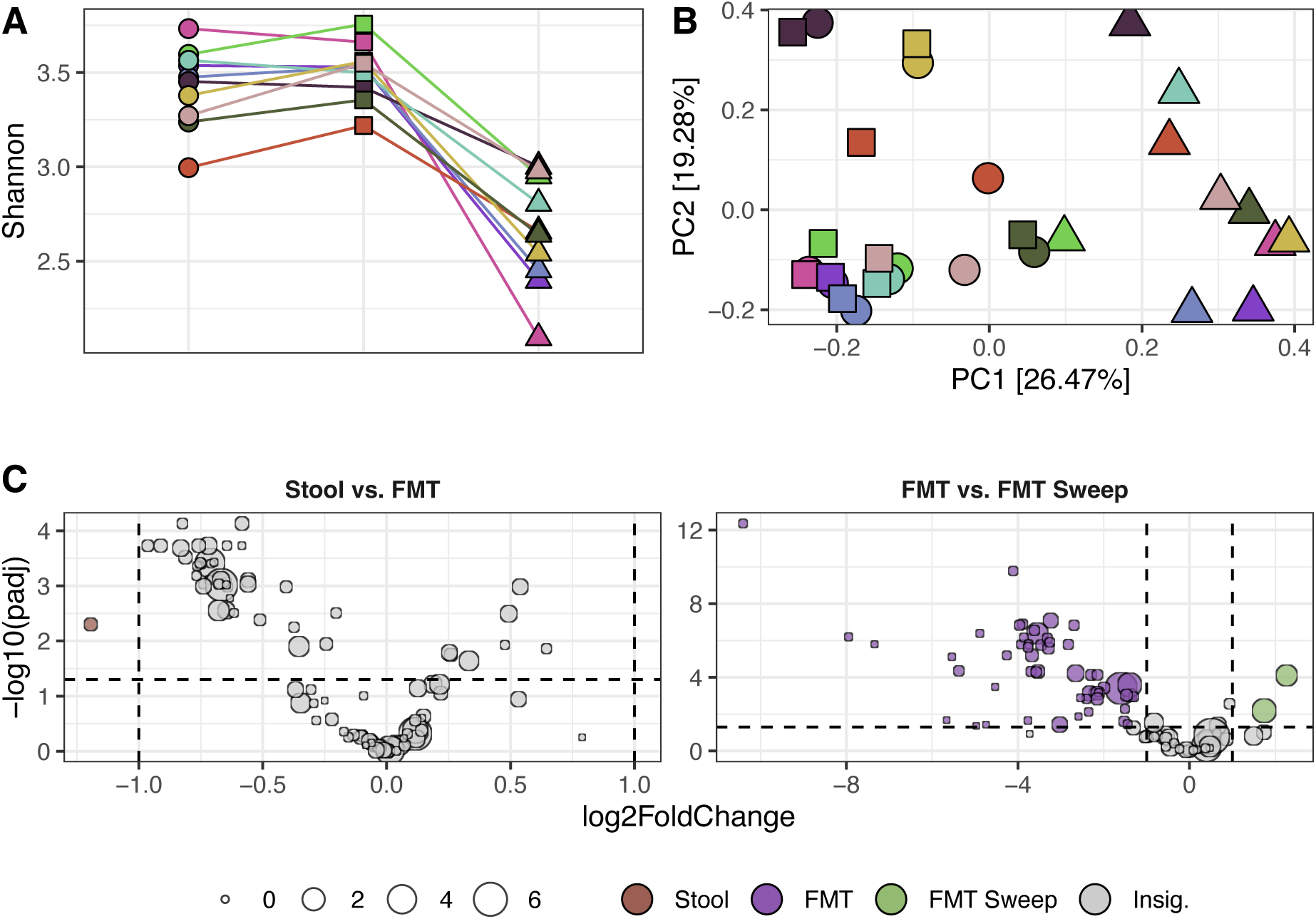
Microbiome Fidelity Between Source Faecal Samples and FMTs. A) Alpha diversity trajectory of microbiome samples by Donation and Source. B) Principal Coordinate Analysis of Bray-Curtis dissimilarities between samples. C) Volcano plot of species differential abundance between sample sources. For panels A and B, each point represents a single microbiome sample; point colour indicates donation and shape indicates source. For panels C, each point represents a single species; point size reflects the median relative abundance of the species across the two sources being compared, and point colour indicates the source in which the species is more abundant. Horizontal dashed lines denote the FDR-adjusted p-value threshold of 0.05, and vertical dashed lines indicate the log₂ fold change cutoff of ±1.

Microbiome composition was significantly influenced by both sample source and donation (PERMANOVA, Bray-Curtis dissimilarity) (Fig. 2B). When all sample sources (Stool, FMT, and FMT Sweep) were included, source explained 25% of the variation (*p* = 0.001), while donation accounted for 53% (*p* = 0.001).

To better understand the contribution of each sample source to overall community variation, separate models were generated excluding one source at a time. In the comparison between Stool and FMT, source explained just 4.2% of the variation (*p* = 0.001), while donation accounted for over 90% (*p* = 0.001), indicating minimal compositional change due to processing relative to between-donor differences. In contrast, comparisons involving FMT Sweep samples revealed much stronger source effects: 27.7% of the variation was explained when comparing FMT to FMT Sweep (*p* = 0.001), and 23.7% when comparing Stool to FMT Sweep (*p* = 0.001). The increased divergence of FMT sweep samples, coupled with reduced Shannon diversity, likely reflects differential growth under culturing conditions, which can selectively enrich for certain taxa based on nutrient availability and growth kinetics. Across all models, donor remained a dominant source of variation, consistently explaining 49-90% of the total community differences.

Pairwise Bray-Curtis dissimilarities were significantly lower between samples from the same donation compared to those from different donations when comparing both Stool vs. FMT (Two-sample Wilcoxon test, *p* = 1.6x10^-^⁶) and FMT vs. FMT Sweep (*p* = 0.005). Intra-donation dissimilarities differed significantly between comparisons (*p* = 1.1x10^-^⁵), with higher divergence observed between FMT and FMT Sweep samples than between Stool and FMT. A similar trend was observed among inter-donation comparisons (*p* = 3x10^-^¹⁸), suggesting that sample source and donor both contribute to variation in microbial composition. Additionally, the repeat donations from the same donor (Donation02 and Donation03) were significantly more similar to each other than to donations from other donors for all three sample sources (Kruskal-Wallis rank sum test, *p* = 0.044 for all) (Table S1).

*Sutterella wadsworthensis* was the only species identified as differentially abundant (|log2FC| ≥ 1, FDR-adjusted *p* ≤ 0.05) between Stool and FMT samples (Fig. 2C). 56 species were identified as differentially abundant between FMT and FMT Sweep samples. *Bifidobacterium longum* and *Collinsella aerofaciens* were more abundant in FMT Sweep samples while 54 species were more abundant in FMT samples. Of these 54 species, the majority (43/54) belonged to the class *Clostridia*; specifically, the *Lachnospiraceae* (20), *Oscillospiraceae* (20), *Peptostreptococcaceae* (2) and *Eubacteriaceae* (1) families (Fig. S1). This taxonomic enrichment pattern was corroborated using *TaxSEA*, which identified consistent overrepresentation of *Clostridia* among species more abundant in FMT samples compared to FMT Sweeps – a pattern that was also evident at the family level which *Lachnospiraceae* and *Oscillospiraceae* consistently depleted in FMT Sweeps (Fig. S2).

*S. wadsworthensis* was a low-abundance species, with a mean relative abundance of 0.44% in Stool metagenomes. In contrast, the 54 species that were significantly more abundant in FMT samples compared to FMT Sweep samples accounted for a substantial portion of the community, with a combined mean relative abundance of 52.5% in FMT metagenomes. The two species enriched in FMT Sweep samples (*B. longum* and *C. aerofaciens*) had a combined mean relative abundance of 3.1%. These results suggest that the culturing process used to generate the FMT Sweep samples selectively enriched a narrow subset of taxa at the expense a broad, but taxonomically-related, group of abundant commensals.

### Microbial Viability is Maintained Post-Processing

To assess whether FMT processing affects microbial viability and recovery, we examined the extent to which microbes present in the original Stool and FMT samples were detectable in the corresponding FMT Sweep. Specifically, we quantified the proportion of total relative abundance and the proportion of species from each source that were recovered. Across donations, 92.9 ± 2.92% of relative abundance and 98.1 ± 1.55% of species were recovered from Stool samples, while 92.1 ± 2.99% of relative abundance and 98.0 ± 1.44% of species were recovered from FMT samples (mean ± SD). Mixed-effects models showed a small significant difference between sample types for relative abundance recovered (estimate = 0. 7104, *p* = 0.031) but not for species count: estimate = 0.1053, *p* = 0.678). These results indicate that processing has a minimal impact on overall viability in FMTs.

Six different species were not recovered by sweep metagenomics in at least one donation. No species were unrecovered in all 10 donations, but three species were unrecovered in a majority of donations. *Ruminococcus sp. FMB-CY1* and the bacteriophage *Carjivirus hominis* were unrecovered in nine donors (all except Donation04), while *Carjivirus communis* was unrecovered in 7 donations.

### Strain Isolation Recovers a Diverse Set of Gut Microbes from FMT Products

Draft genome assemblies were generated for 24 strains isolated from each FMT. Following genome assembly and quality control, a total of 191 high-quality genomes were retained, with a median of 20 genomes per donation (range: 13–23). Of these, 176 genomes were classified to species level, representing 65 distinct species spanning five bacterial phyla: *Actinobacteriota*, *Bacteroidota*, *Firmicutes*, Firmicutes_A, and *Firmicutes_C* (Fig. 3A). The 15 unclassified genomes were all identified as members of the *Collinsella* genus.

**Figure 3.**
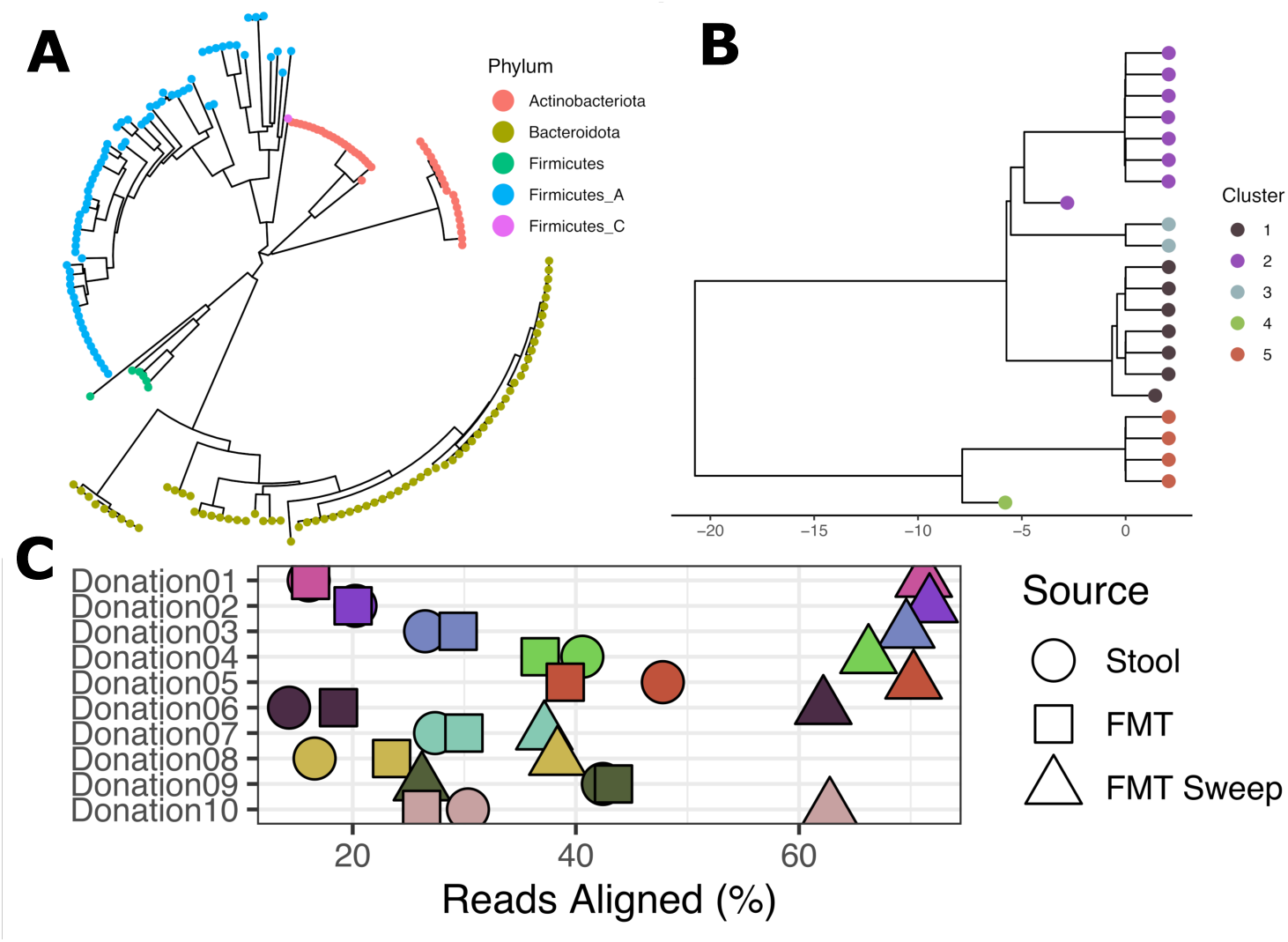
Recovery of viable microbes from FMT samples using strain isolation and sweep metagenomics. A) GTDB-tk-derived phylogenetic tree of all high-quality isolates (*n* = 191) rooted around the *Actinobacteriota* phylum. Tip points are coloured by phylum. B) A dendrogram of *Collinsella* genomes generated using complete-linkage hierarchical clustering of pairwise ANI distances, expressed as (100 – ANI%). Tip points are coloured by species-level cluster membership, defined using a 95% ANI threshold. Cluster 1 contains only genomes classified as *Collinsella aerofaciens_J*, while all remaining clusters represent putatively novel or unclassified *Collinsella* species. C) Proportion of the metagenome recovered by the isolated strains, based on alignment of metagenomic reads from Stool, FMT, and FMT Sweep samples to genomes of isolates from the paired FMT sample. Each point represents a sample; shape indicates sample source, and colour indicates donation ID.

**Figure 4.**
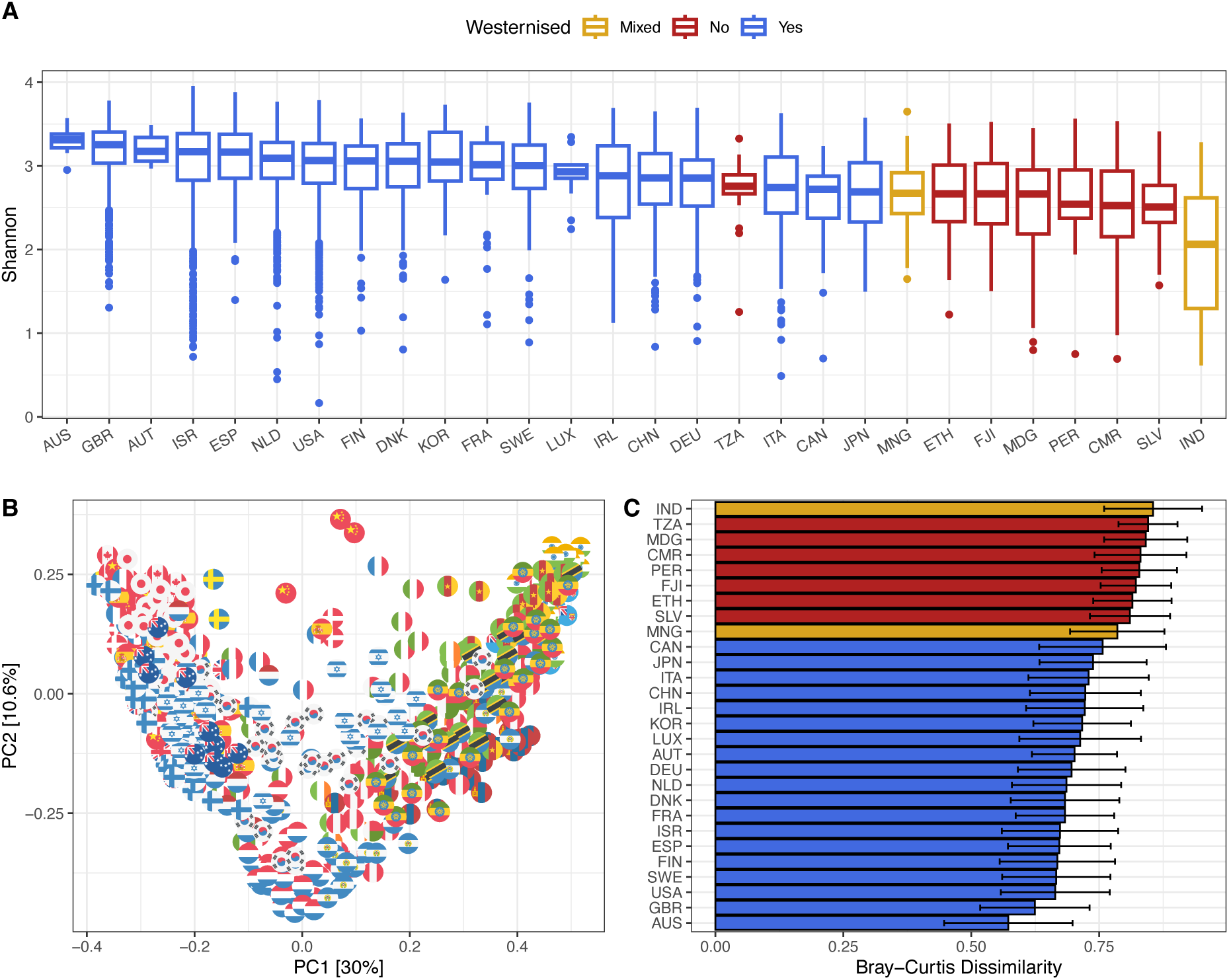
Comparison of Australian FMT donor microbiomes to global healthy stool microbiomes. A) Shannon diversity of stool metagenomes stratified by country of origin and ordered by median Shannon diversity. B) Principal Coordinate Analysis of Bray-Curtis dissimilarities between samples from different countries. Up to 30 representative samples per country are shown for ease of interpretation, selected based on the lowest median pairwise Bray-Curtis dissimilarity to other samples within the same country. C) Distribution of Bray-Curtis dissimilarities to Australian samples stratified by country and ordered by mean dissimilarity.

The most frequently isolated species were *Blautia_A wexlerae* (14 isolates from 7 donations), *Phocaeicola vulgatus* (12 isolates from 7 donors), *Bifidobacterium longum* (9 isolates from 6 donations), *Bariatricus comes* (7 isolates from 5 donations), and *Mediterraneibacter faecis* (7 isolates from 5 donations).

Relative evolutionary divergence (RED) values ranging from 0.935 to 0.997 suggest the 15 unclassified *Collinsella* isolates likely belong to one or more novel or currently undescribed species within the genus. Pairwise ANI comparisons, combined with complete-linkage hierarchical clustering and species-level thresholds (ANI ≥ 95%), suggest that these isolates represent four distinct species, bringing the total number of *Collinsella* species isolated in this study to five (Fig. 3B).

To evaluate the representativeness of the strains isolated from each FMT sample, metagenomic reads from matched Stool, FMT, and FMT Sweep samples were aligned to genome assemblies of the corresponding cultured isolates. The proportion of reads aligning to these genomes did not differ between Stool (mean: 28.2%) and FMT (mean: 28.3%) samples (estimate = -0.112, *p* = 0.9998). In contrast, alignment rates were significantly higher in FMT Sweep samples (mean: 57.6%) compared to both Stool (estimate = -29.338, *p* = 0.0002) and FMT (estimate = -29.226, *p* = 0.0002), consistent with likely selective enrichment of culturable taxa under the sweep culturomics conditions used for strain isolation (Fig. 3C).

No strains from the phylum *Pseudomonadota* (formerly *Proteobacteria*) were isolated, likely reflecting their low abundance in the gut microbiome. This is consistent with the metagenomic data, in which the mean *Pseudomonadota* relative abundance was 1.07%, in line with levels reported in healthy adults^43^. There was no significant difference in *Pseudomonadota* relative abundance in FMT (estimate = -0.1692, *p* = 0.7739) or FMT Sweep (estimate = -0.4262. *p* = 0.472) samples compared to source Stool samples (Fig. S3).

### Detection of Strain-Sharing Across Sample Sources Varies by Species

Strain-resolved metagenomic analysis revealed that individual strains were traceable across stool, FMT, and FMT Sweep samples; however, the number of detectable strain clusters and the extent of cross-sample sharing varied markedly by species. *Blautia_A wexlerae* and *Bacteroides uniformis* exhibited the highest levels of intra-donation strain sharing, with 14 and 10 strain clusters, respectively, detected in more than one sample from the same donation (Figure S4). Notably, Donation05 contained six distinct *B. wexlerae* strain clusters with evidence of sharing across sample types. The repeat donations from the same donor (Donation02 and Donation03) shared more strains than donations from independent donors for all three sample sources (Table S2). This variation in detectable strain sharing likely stems from a combination of factors, including species-specific abundance (which affects detection sensitivity), sequencing depth (with deeper sequencing potentially revealing additional shared strains), and differences in strain resilience to processing.

### Australian FMT Donors Show High Diversity and Regional Similarity to Western Cohorts

To understand how the faecal microbiome of healthy Australian FMT donors compares to those from other populations, we compared our source Stool samples to 7,014 stool samples from the *curated MetagenomicData* R package, representing 53 studies conducted across 27 countries (Supplementary material).

Shannon diversity was significantly higher in Australian samples (source Stool samples in this study) compared to all other countries except Austria (*p* = 0.670), Spain (*p* = 0.121), the United Kingdom (*p* = 0.345), and South Korea (*p* = 0.130), where the differences were not statistically significant. The most pronounced differences were observed in samples from India (estimate = -1.317, *p* = 1.07x10^-17^), followed by Madagascar (estimate = -0.8, *p* = 1.1e-7) and Cameroon (estimate = -0.791, *p* = 1.34x10^-7^), suggesting substantial differences in microbial richness and evenness across geographic regions.

Microbiome profiles from other countries were significantly more dissimilar to Australian samples than intra-Australian comparisons (mean dissimilarity: 0.572). The most similar microbiomes to Australian samples were from the United Kingdom (estimate = +0.052, *p* = 0.037), the United States (estimate = +0.088), and Sweden (estimate = +0.093), while the most distinct were from India (estimate = +0.283), Tanzania (estimate = +0.273), Madagascar (estimate = +0.269), and Peru (estimate = +0.259) (all *p* < 0.001 unless stated). These findings highlight both global microbiome diversity and the influence of geographic and lifestyle factors on community composition.

## Discussion

This study provides a detailed characterisation of faecal microbiota transplant (FMT) samples from volunteer healthy Australian donors through integrated metagenomic and culturomic analyses, offering new insights into the consistency of donor microbiomes, the fidelity of FMT processing, and the viability of recovered microbial communities.

Longitudinal analysis of repeat donations over a period of up to 55 days demonstrated high temporal consistency in the microbiome profiles of individual donors, reinforcing the reproducibility and reliability of sourcing FMTs from repeat donations by the same individual and supporting the current practice of banking multiple samples from approved donors. Temporal stability is particularly important for ensuring standardised FMT composition across different treatment batches and timepoints, a critical consideration for both clinical efficacy and regulatory oversight. However, the strong donor-specific signatures observed also suggest that treatment outcomes may vary depending on the microbial composition of the donor. In cases of treatment failure or non-response, re-treatment with FMTs derived from a different donor may improve efficacy by providing a more compatible or therapeutically active microbial community.

Microbiome comparisons between source faecal samples and corresponding FMT preparations revealed minimal differences in community structure, diversity, and taxonomic composition. While statistically significant differences were detectable, these were minor and largely overshadowed by inter-donation variability. This high level of compositional fidelity suggests that FMTs preserve the essential microbial features of the donor material and are not substantially altered by processing in ways that could affect their therapeutic potential. It also reinforces the value of robust donor screening, as donor-specific features are likely to play a major role in treatment outcomes^12^.

The donor-specific microbiome signatures described here highlight the need to consider both individual donors and individual donations when defining what constitutes an “optimal” FMT product. While all donors in this study passed rigorous screening protocols, their microbiome profiles were relatively heterogeneous. Incorporating metagenomic profiling into the donor screening process may help guide the selection of an alternative donor or donation in cases of treatment failure and, as our understanding of the microbial traits associated with FMT success continues to grow, this information can be used to prioritise FMT products with the highest predicted likelihood of clinical benefit.

Sweep metagenomics and strain isolation demonstrated that a high proportion of the microbial community, both in terms of abundance and species richness, was recoverable from stool and FMT samples. These results highlight the potential of FMTs to transfer a viable and functionally relevant microbial community capable of colonising the recipient gut.

FMT Sweep samples showed significantly higher alignment to the genomes of isolated strains, likely reflecting culturing biases that selectively enriched for taxa amenable to growth under YCFA-based conditions. Of the 54 species found to be less abundant in FMT Sweep samples compared to their corresponding FMTs, 43 belonged to the *Clostridia* - a class rich in spore-forming taxa^44^. This underrepresentation is consistent with the known resistance of spores to standard YCFA culturing conditions in the absence of ethanol-based enrichment methods, which were not employed in this study^45^. Coupled with the high proportion of the microbiome deemed recoverable, this suggests that variation introduced by culturing conditions accounts for the majority of observed differences, with FMT processing itself contributing minimally to compositional change.

When contextualised against publicly available datasets spanning 27 countries, the faecal microbiomes of healthy Australian FMT donors aligned most closely with microbiome profiles from other Western populations. Shannon diversity was significantly higher in Australian samples compared to most countries analysed, except for a few Western nations (Austria, Spain, the United Kingdom) and South Korea, where differences were not statistically significant. Microbiome compositions from Australia were most like those from the United Kingdom, the Netherlands, and Spain, while the largest dissimilarities were observed with samples from India, Madagascar, Cameroon, and Peru. These differences underscore the substantial geographic variation in gut microbiota and highlight the importance of considering population context when selecting FMT donors.

These insights also have implications for the development of standardised donor selection protocols across diverse populations. FMTs derived from donors with westernised microbiome profiles may not be universally representative or appropriate for use in all populations, particularly in settings with markedly different microbiome baselines, and determinants of FMT efficacy may differ between populations. Future efforts should explore whether tailoring donor selection to match recipient microbiome backgrounds improves engraftment and clinical outcomes, particularly in diverse settings.

While this study provides a comprehensive characterisation of FMTs using metagenomic and culturomic approaches, several limitations should be acknowledged. First, the sample size and donor diversity were limited, potentially constraining the generalisability of findings across broader populations. Expanding donor cohorts with greater variation in age, diet, ethnicity, and microbiome composition will be important for assessing the robustness of FMT characteristics across diverse clinical contexts. Second, the culturing approach used - based solely on YCFA - likely introduced selection biases, favouring taxa that are fast-growing or metabolically adapted to the specific nutrient conditions provided. This bias may underestimate the viability of more fastidious, functionally important anaerobes. Future work incorporating broader culturing strategies may capture a more representative set of viable community members. Finally, while comparisons to global public datasets were conducted using the same software and database versions, the nature of publicly available data means that some degree of technical variation - particularly from upstream sample handling and sequencing protocols - cannot be fully excluded and should be considered when interpreting inter-cohort differences.

Our findings also have potential regulatory implications. The observed consistency across repeat donations and between source and processed FMTs supports the suitability of the employed donor screening and manufacturing pipelines, which are critical for reproducibility, safety, and regulatory approval. These data could inform quality control criteria for FMT product development and help establish benchmarks for therapeutic consistency and viability.

## Availability of data and materials

Sequencing data generated in this study are available at BioProject PRJNA1303742. The metagenomic data have been uploaded following quality trimming and removal of human DNA to ensure that host genetic data is not made publicly available.

---

**Figure S1.**
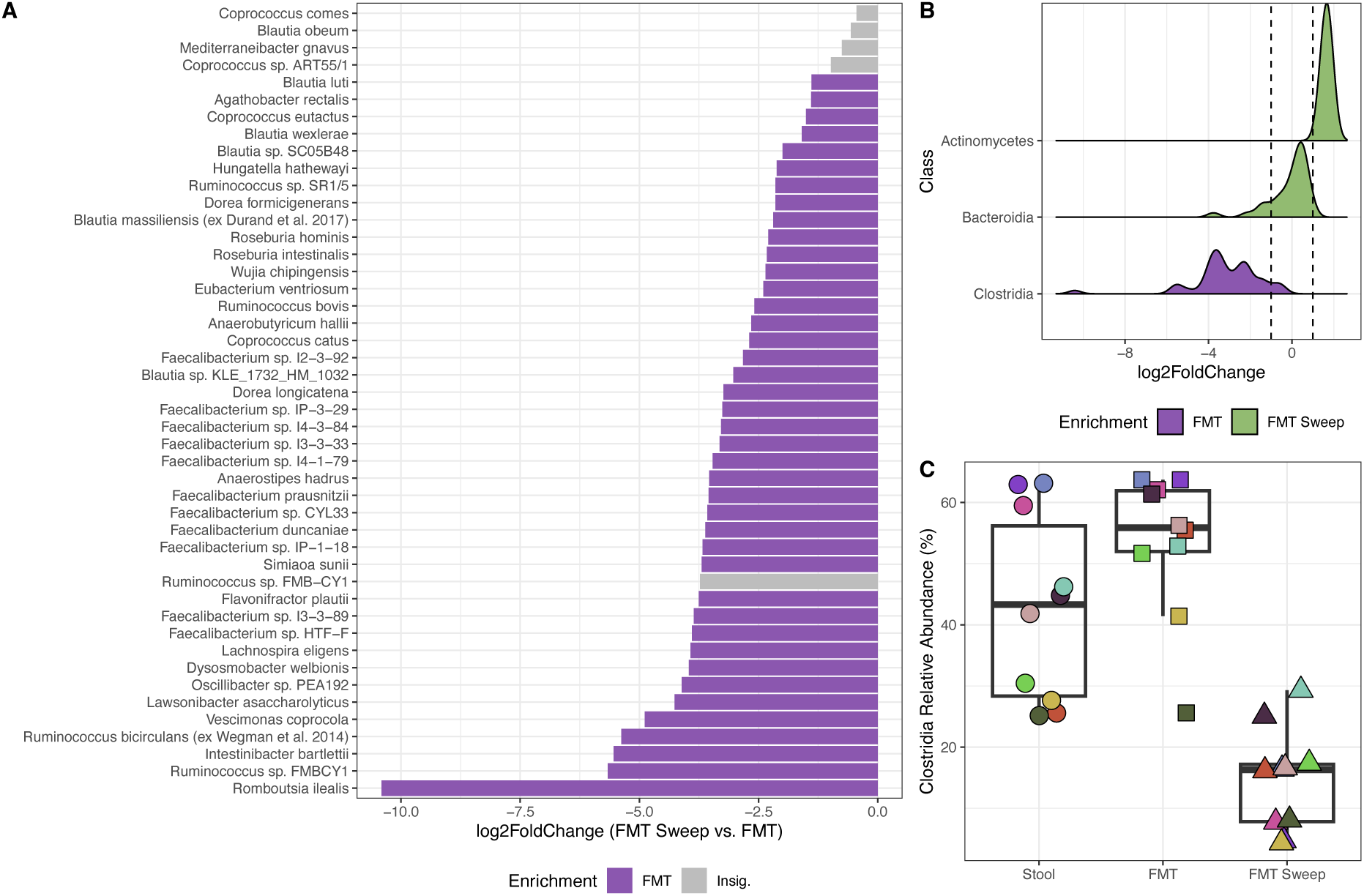
Impact of YCFA sweep metagenomics on *Clostridia* species. A) Differential abundance of species in the *Clostridia* class as measured by *LinDA* when comparing FMT Sweep samples to their corresponding FMT sample. B) Distribution of species log2 fold changes grouped at the class level of taxonomy. Only classes determined to be significantly enriched by *TaxSEA* are shown and density plots are filled by sample source in which the class is enriched. Vertical dashed lines represent log2FC values of -1 and +1. C) Summed relative abundance of all *Clostridia* species in each sample, stratified by sample source. Points are coloured by donation ID.

**Figure S2.**
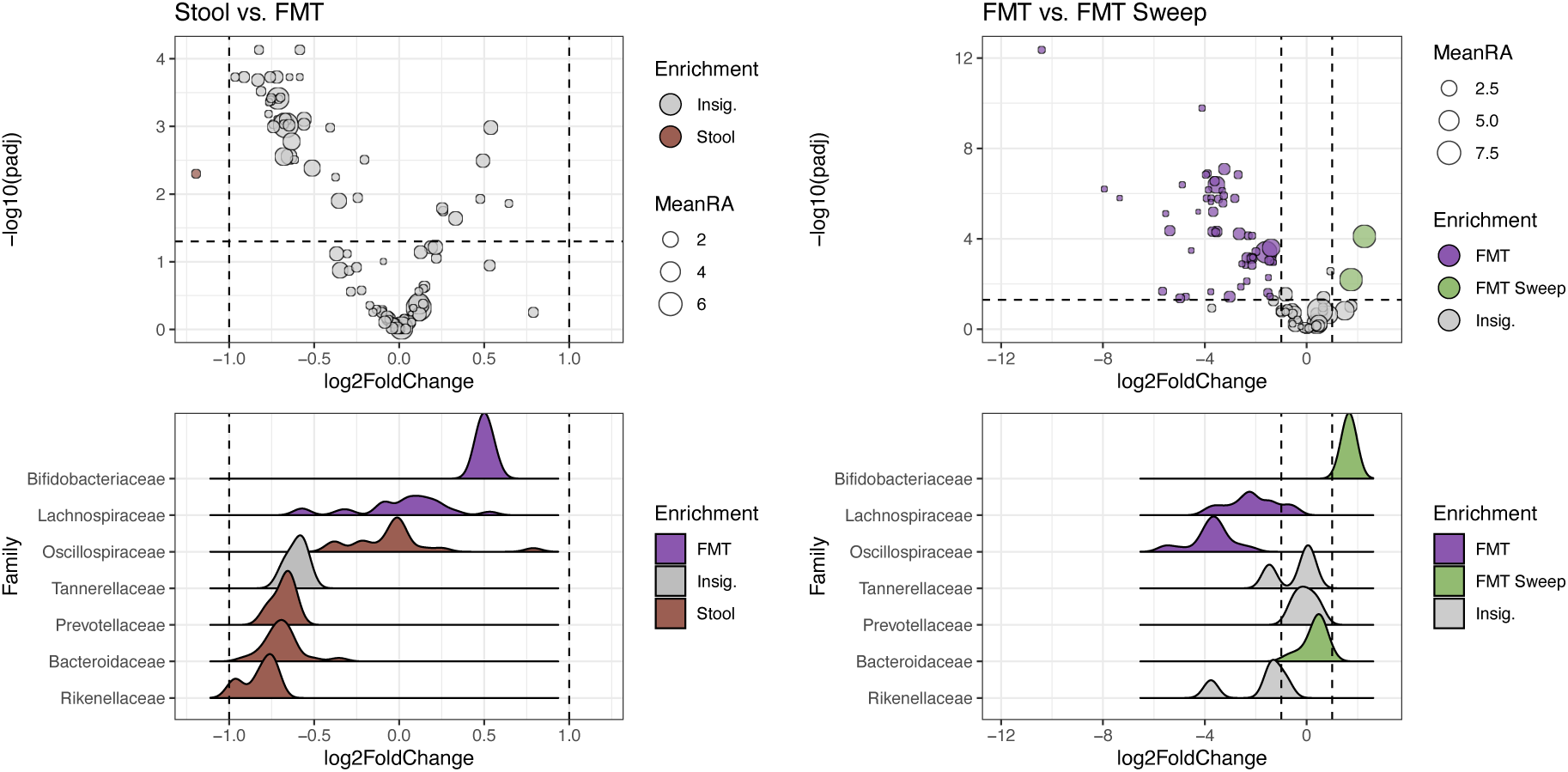
Impact of FMT processing and YCFA sweep metagenomics on microbiome composition at the species and family level. Top row: Species-level differential abundance results as measured by *LinDA* when comparing FMT samples to their corresponding Stool sample (left), and FMT Sweep samples to their corresponding FMT sample (right). Bottom row: Distribution of species log2 fold changes grouped at the family level of taxonomy in each comparison. Only families with 3 or more species were included in *TaxSEA* analysis and density plots are filled by sample source in which the family is enriched. Vertical dashed lines represent log2FC values of -1 and +1.

**Figure S3.**
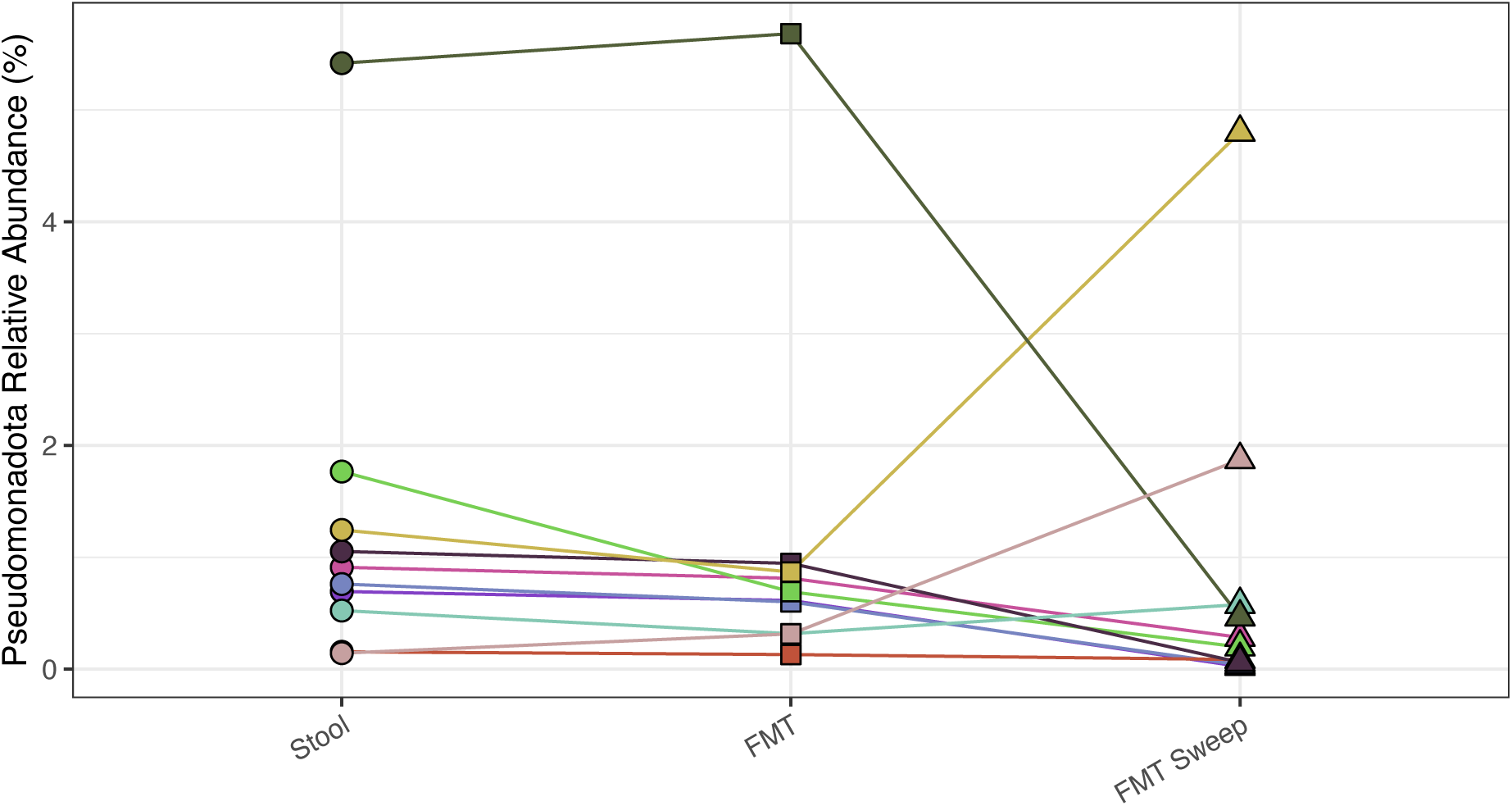
*Pseudomonadota* relative abundance across sample sources. Phylum-level relative abundance calculated by Kraken2 and Bracken. Each point represents a sample, shape denotes source, and colour denotes donation ID. Lines are added to allow easier interpretation of relative abundance changes between sample sources from the same donation.

**Figure S4.**
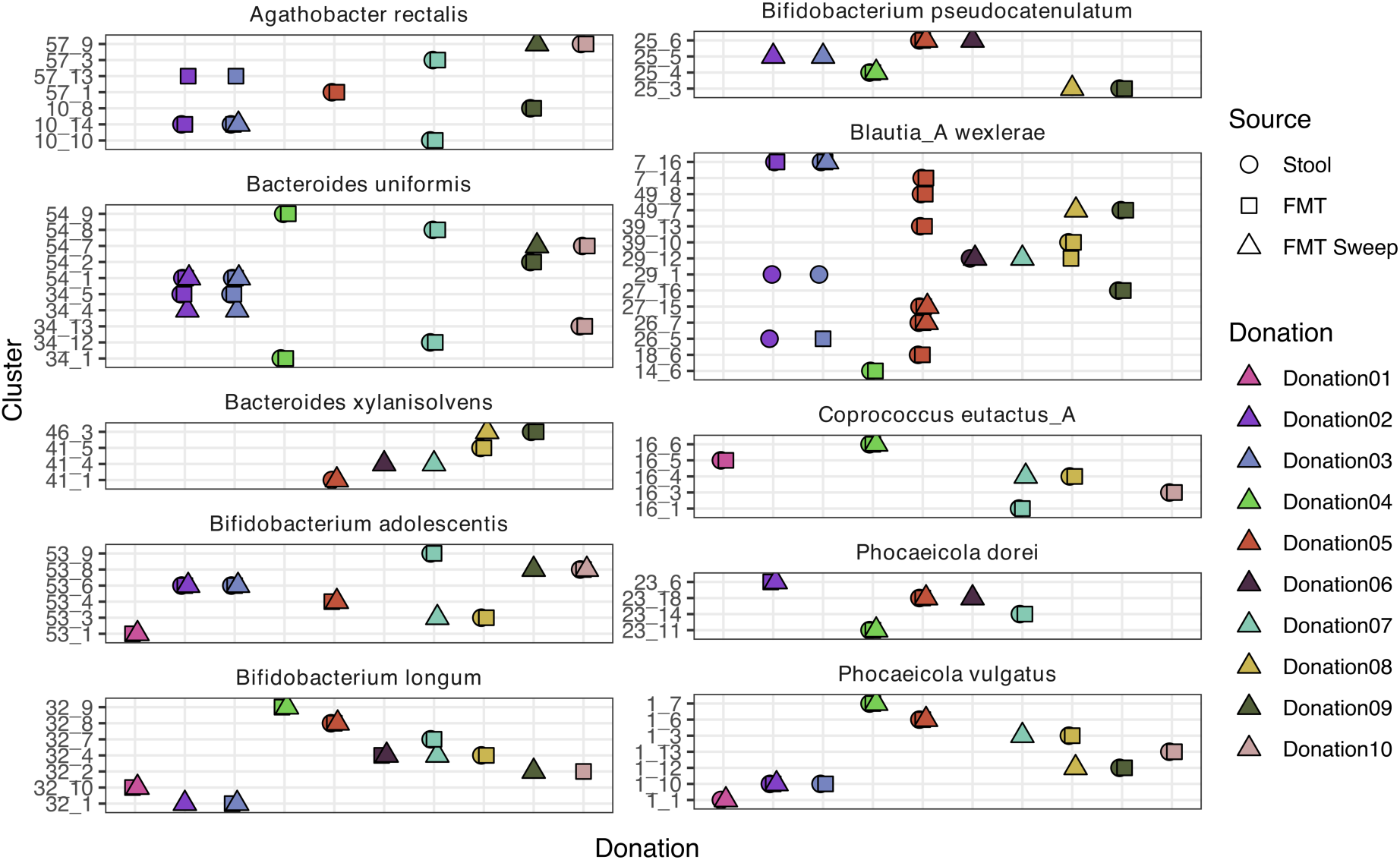
Strain cluster co-occurrence across donations and sample sources. Strain-level co-occurrence patterns for species with evidence of strain sharing between samples, as identified by *inStrain*. Each facet displays results for a single species. Species were ranked by the number of strain clusters with only the top ten shown. Rows represent distinct strain clusters, and points indicate the presence of that strain cluster in a given metagenomic sample. Point shape denotes the sample source (Stool, FMT, or FMT Sweep), and colour corresponds to donation ID.

**Table S1.** Bray Curtis dissimilarities of repeat donations from the same donor compared to pairwise distances between donations from different donors.

| RepeatDonor | Source | Bray-Curtis (Mean) | Bray-Curtis (SD) |
| --- | --- | --- | --- |
| FALSE | Stool | 0.54 | 0.113 |
| TRUE | Stool | 0.132 | - |
| FALSE | FMT | 0.494 | 0.102 |
| TRUE | FMT | 0.0976 | - |
| FALSE | FMT Sweep | 0.578 | 0.112 |
| TRUE | FMT Sweep | 0.203 | - |

**Table S2.**
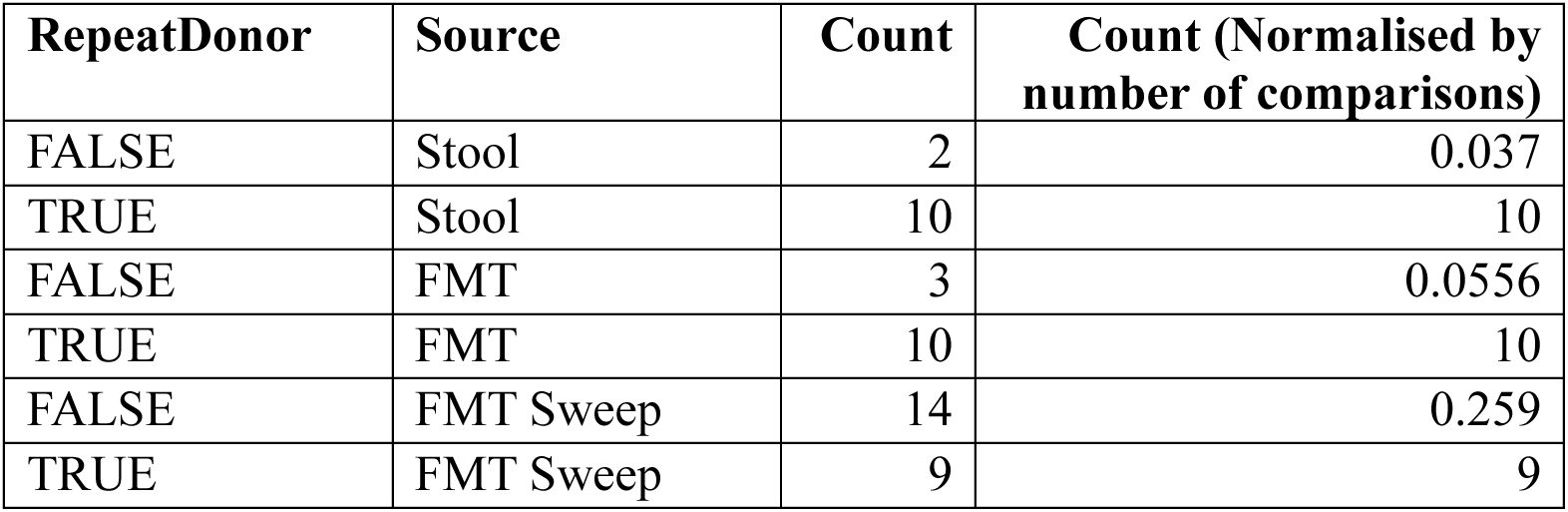
Strain sharing events detected in repeat donations from the same donor compared to donations from different donors.

| RepeatDonor | Source | Count | Count (Normalised by number of comparisons) |
| --- | --- | --- | --- |
| FALSE | Stool | 2 | 0.037 |
| TRUE | Stool | 10 | 10 |
| FALSE | FMT | 3 | 0.0556 |
| TRUE | FMT | 10 | 10 |
| FALSE | FMT Sweep | 14 | 0.259 |
| TRUE | FMT Sweep | 9 | 9 |

